# Abundance and Distribution of *Heterelmis cf. glabra* (Coleoptera: Elmidae) within Dolan Falls Preserve and the Devils River State Natural Area, Texas, USA

**DOI:** 10.1101/2020.02.10.941641

**Authors:** Peter H. Diaz, J. Randy Gibson, Chad W. Norris, Carrie L. Thompson

## Abstract

The Devils River watershed in south-central Texas has baseflows entirely attributable to groundwater primarily sourced from the Edwards-Trinity and Edwards Aquifers. The largest known populations of a species of riffle beetle, *Heterelmis cf. glabra*, are located in springs associated with the upper Devils River. The focus of this study was to 1) determine site-level abundances of *H. cf. glabra* using open system N-mixture models, 2) test mesohabitat associations of members in the riffle beetle family Elmidae and, 3) measure and examine abiotic and habitat associations for adult and larval beetles within the study area. We sampled 32 spring sources to determine occupancy and abundance of adult and larval riffle beetles (Elmidae) within the study area. Spring sources were mapped and categorized by type (orifice, upwelling, group of springs, or seep). Basic water chemistry and flow rate categorization were also performed at each site. Model results suggest that rainfall, flow and site are important for detection of *H. cf. glabra*. Based on our results, regular monitoring of these 32 sites using these methods, is recommended to conduct hypotheses tests on covariates influencing abundance. Such baseline information will be important in measuring impacts to this and other spring-associated species as the habitats of this region are impacted by natural or anthropogenic phenomena.

## Introduction

Groundwater extraction in the Permian Basin of West Texas has increased as industrial pumping for gas and oil have increased. In addition, the region has been identified as a potential source for water export to more highly populated regions of Texas [1]. The depletion of groundwater in certain areas may in turn cause disruptions in flow or changes to historically stable temperatures that endemic spring-adapted species have adapted to over the course of geologic history. Many spring-adapted species are known to be associated with stenothermal groundwater habitats of the Edwards Plateau of Central and West Texas [2–6]. Although springs may fluctuate in regards to flows over geological periods of time, direct correlations between flows and anthropogenic pressures are observable in nearly real time in unconfined aquifers such as the Edwards-Trinity aquifer that feed the Devils River [1]. This in turn may alter the characteristics of the spring system under which these species naturally persist. Recent modeling suggests that the impact of groundwater withdrawals on Devils River discharge is proportional to the amount of water pumped [1]. Subsequently, significant groundwater pumping has the potential to extinguish or shift the location of spring discharge points. During times of drought or disturbance, some spring-adapted species are able to retreat into the aquifer for temporary refuge [7] or live deeper within the aquifer permanently [8]. Other spring adapted-organisms, such as *Heterelmis comalensis* and *H. glabra*, have life history patterns requiring surface components, which makes them more susceptible to changes in springflow that alter the surface habitats condition.

*Heterelmis cf. glabra* represents a potentially undescribed species of riffle beetle, is known to exist in large permanent springs in Terrell, Val Verde, Kerr, Hays, Bell, and Tom Green counties [9,10, unpublished data]. [9,10]. Despite recent applied research evaluating tolerance to elevated temperatures and reduced dissolved oxygen for a population of *H. cf. glabra* [11, 12], little information is available on the distribution and abundance of this species within its known range or its habitat associations. Gathering and analyzing reliable data on population size and distribution is a central theme in ecological research [13] and is essential for management of endemic populations [14].

The Comal Springs riffle beetle (*Heterelmis comalensis*; [15]) is a USA federally endangered species [16] hypothesized to be similar to *H. cf. glabra* both phylogenetically [9, 10] and ecologically (i.e., spring obligate). The major threats to *H. comalensis* are reduction in water quality and quantity due to drought and development associated with an increased need for groundwater resources as a result of accelerated population growth in the area [17, 18]. Similar to *H. cf. glabra*, *H. comalensis* inhabits areas near and within spring sources [19, 8] and are often found associated with woody debris or roots where they feed on biofilm produced as these substrates decay [20, 8, 18, 21]. *Heterelmis comalensis* are thought to move through interstitial alluvium within spring sources, making collections difficult as much of this habitat is not accessible by traditional sampling techniques. Subsequently, a method was developed using cotton lures placed in spring sources for monitoring *H. comalensis* [22]. Over several weeks, biofilms on which riffle beetles feed grow on the cotton material. This provides a consistent method of collection for this endangered species. Using this method, hundreds of larvae and adults can be collected and returned to the habitat unharmed [8, 23]. Populations of *Heterelmis cf. glabra* in large perennial springs of the Edwards Plateau occupy ecologically similar habitat as *H. comalensis* and are readily collected using the cotton lure methodology.

The life history characteristics of these riffle beetles provide statistical and study design challenges for population monitoring. Adult *Heterelmis* beetles probably live about a year (San Marcos Aquatic Resources Center - unpublished captive propagation data) and are small (~2 mm) creating issues with mark recapture studies [23]. Certain interstitial spring-adapted species most likely occupy areas within spring sources not accessible to sampling gear such as a Hess sampler or kick net producing low count data not or with many zeros. The use of count data for a level of abundance without taking into account the organisms not detected can be misleading, by invoking a suspect relationship between the count index and true abundance [24]. To rectify this discrepancy, advances in monitoring techniques can allow for estimation of abundance using count data and covariates that partition the distribution of the target organism spatially while accounting for imperfect detection [25, 26]. These models are called N-mixture models and are a class of state space models which assume the system is observed imperfectly [27]. These models can be used on open [26] or closed systems [25].

The focus of this study was to determine site-level abundances of *H. cf. glabra* within the study area. This was accomplished by testing a series of models based on hypotheses associated with the detection of the beetle. Other objectives of the study include testing spring associated affinities of members in the riffle beetle family Elmidae present within the system. Measured abiotic associations and basic habitat associations for adult and larval beetles were examined.

## Methods

### Study Area

The Devils River watershed is in south-central Texas and is one of two principal Texas tributaries to the Rio Grande. The Devils River is primarily sourced by the Edwards-Trinity Aquifer, with baseflows entirely attributable to groundwater [1]. The largest known populations of *H. cf. glabra* are located in Finegan, Blue, and Dolan springs. These springs issue into the upper Devils River portion of The Nature Conservancy’s Dolan Falls Preserve (DFP) and Texas Parks and Wildlife Department’s Devils River State Natural Area (DRSNA) property (Fig 1).

**Fig 1.**
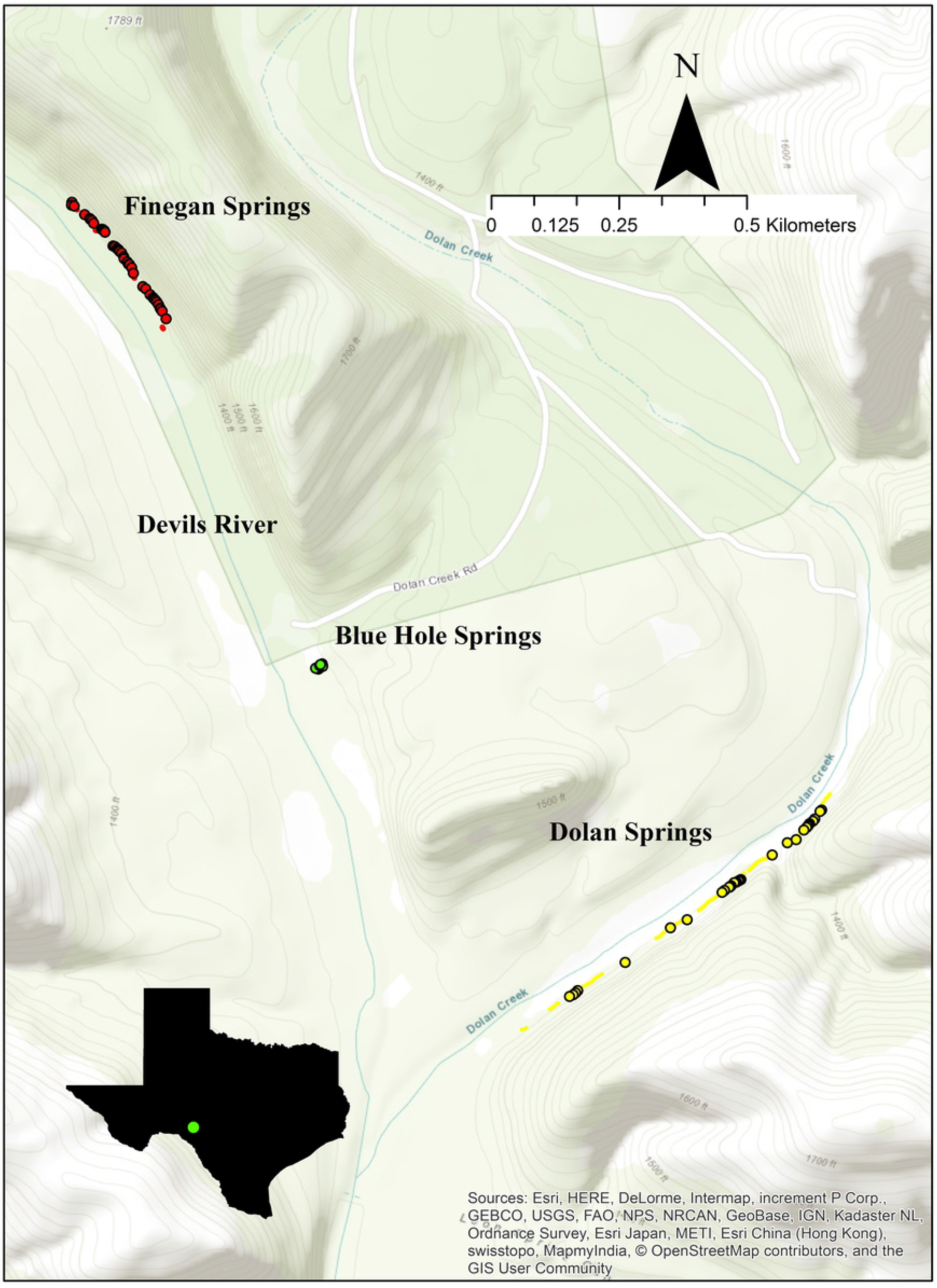
Map of study area in Val Verde County Texas. Mapped springs on The Nature Conservancy’s Dolan Falls Preserve and Texas Parks and Wildlife Department’s Devils River State Natural Area. Sites were selected randomly from these available springs.

The spring complexes in the study area issue from Cretaceous Edwards Limestone [28] along the east bank of the Devils River and Dolan Creek. Finegan Springs comprises (discharge = 99-760 L/s from 5 measurements during 1928-1971; [29]) 44 mapped springs and seeps along a stretch of around 333 m at the base of a bluff flowing over chert bedrock forming small rheocrene streams that merge into larger pools and streams emptying into the Devils River as far as 25 m from spring sources. Blue Springs is a small group of spring conduits and gravel seeps with most (5 springs/seeps) flowing into a short (ca. 5 m) cobble/gravel rheocrene and four marginal seeps that empty into a backwater pool (ca. 90 x 30 m) that connects directly to the Devils River ca. 800 m downstream of Finegan Springs. Dolan Springs comprises (discharge = 34-510 L/s from 7 measurements during 1928-1970; [29]) 48 springs or seeps along a stretch of 762 m at the base of a bluff and flow over limestone bedrock forming small rheocrene streams and shallow pools that empty into Dolan Creek as far as 30 m from spring sources. The confluence of Dolan Creek with the Devils River is ca. 500 m downstream of Dolan Creek from the stretch of Dolan Springs and is ca. 1 km downstream of the Devils River from Blue Springs (Fig 1). Water quality issuing from 12 spring sites (seven from Finegan; two from Blue, and two from Dolan springs) in February 2010 (average temperature ≈ 22 °C; conductivity ≈ 504 μS/cm; pH ≈ 7.1; dissolved oxygen ≈ 7.9) were similar to those measured during this study from 64 spring sites in February 2016 (average temperature ≈ 22 °C; conductivity ≈ 494 μS/cm; pH ≈ 7.2; dissolved oxygen ≈ 7.9). These springs are habitat for several rare endemic stygobiontic species including insects, crustaceans, and salamanders [30, 2].

### Data collection and N-mixture model

Individual spring sources were mapped during the week of January 12, 2016. Data collected during the mapping event consisted of basic water chemistry (temperature, dissolved oxygen, pH, conductivity, and total dissolved solids) and a categorical designation of flow from one to five (five being the highest). Springs were identified and categorized as orifice, upwellings, group of springs, and seeps. The designation, “groups of springs”, was used when the springs were too close in proximity to each other to allow the Global Positioning System (GPS) unit to distinguish between the individual orifices accurately. Different types of springs and their placement within the system (above or below the waterline) may have effects on the types of invertebrate communities present. For this study, all of the mapped locations had sites above the waterline. Therefore, sites were randomly selected from two groups within the mapping data (groups of springs/orifice and seeps). Fourteen sites were selected from Dolan, 14 sites from Finegan and four sites from Blue springs. Although seeps were the second most abundant spring type available, most consisted of a thin layer of water moving over bedrock which is not conducive for the cotton lure sampling method as it requires water depths of at least 2 cm. Within the 32 sites, six seeps were selected randomly from the data set although not all were used for previously mentioned reasons. Sites are, at a minimum, a meter apart or separated by terrestrial environment to maintain independence.

To examine abundance of *H. cf. glabra* within the study area, lures were deployed in February, May, August and November of 2016. Each event consisted of burying a folded cotton cloth encased in a metal cage in the substrate of the spring source outflow (Fig 2). The cotton cloth lure is standardized in size (15 cm x 15 cm) and folded the same for each cage. The cage is used to hold the shape of the lure over time and prevent potential anoxia by becoming squeezed [23]. Thirty two lures were set in springs for each of the four events. Each lure represents a sampling site and were left in the substrate for 28 to 35 days to allow for biofilm growth. After that time, the lures were removed and adult and larval riffle beetles (Elmidae) found on each lure were counted and recorded. All riffle beetles were carefully returned to the site of capture following each event. For each subsequent event, this process was repeated at the same sites.

**Fig 2.**
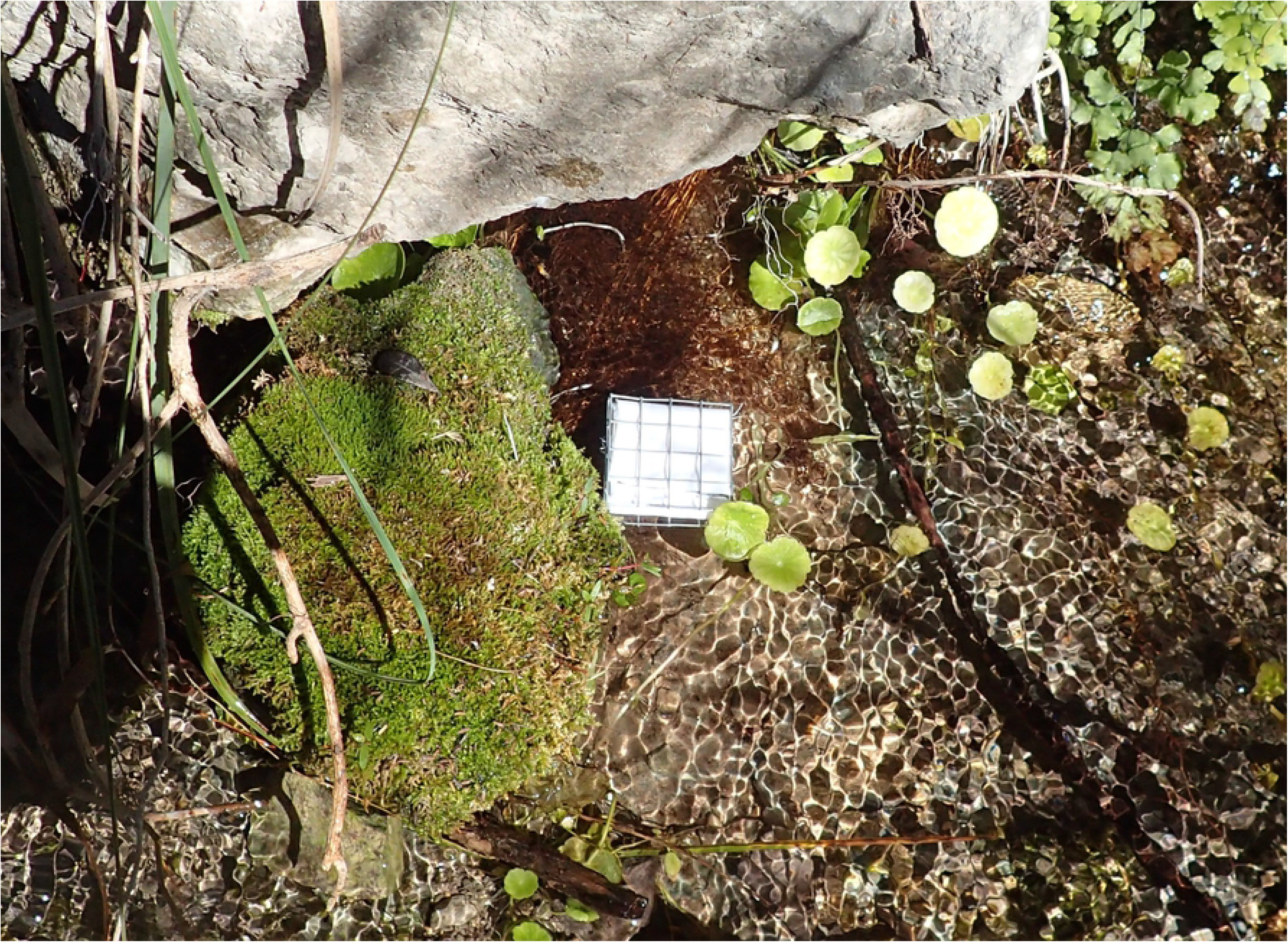
Cotton lure. Cotton lure and cage used to collect riffle beetles to determine site level population estimates from springs in Val Verde County TX.

Since the abundance models use probability of detection in determining the organism abundance, covariates that may influence detection of *H. cf. glabra* were examined. Spring location was considered a site covariate and consisted of grouping sites on the Devils River (Finegan and Blue springs) as one site and Dolan Springs as the other. Flow was considered an ordinal site covariate and ranged from one to five (five having the most flow). Sites with a flow of one were considered seep sites, and ranged from wet rock to a small pooling area with organic debris. Sites with flows above one would be considered more traditional springs with a run terminating at the creek or the river.

Most of the springs arise from the bottom of bluffs that run parallel to the creek or river. During rain events, material is dislodged from higher elevations along the bluff and deposited over the spring sites. This may cause lures to be covered by silt, reducing dissolved oxygen and thereby decreasing the available area for food resources and beetle respiration thus influencing the detection of the beetles after rain events. In addition, rain can cause changes in water chemistry at a local level potentially influencing the areas where riffle beetles would associate. To account for this, rain events during the sampling period were considered as a sampling covariate which has an associated binary score for each sampling event. A score of 1 was given to the model for rain if the rainfall total for a day exceeded 2.54 cm, examined over the duration while the lure was deployed. Rainfall totals were determined by checking the Del Rio International Airport station accessed through Weather Underground (www.wunderground.com).

Capture data of adult, and not larval, *H. cf. glabra* from each event was used to populate the model. The software package “unmarked” in R [31, 32] was used to analyze the data for abundance using the function “pcountOpen” for fitting the open population models [26]. Open models were run due to the relatively short life span of the beetles and the duration of the study. Lost lures during an event were accommodated as per the unmarked manual within the model framework. Model selection was aided using Akaike’s information criterion corrected (AICc) for small sample size [33]. To examine the fit of the model, goodness of fit test were conducted in the R package “AICcmodavg” with 100 parametric bootstrap iterations. The final models were selected based upon the AICc score and the goodness of fit tests.

Candidate abundance models were analyzed using two different abundance distributions: a Poisson distribution with K set at 200 and negative binomial (NB) models with K set at 400. Abundance can vary based on the selection of K (upper bounds of integration), especially when low detection probabilities are calculated [34]. Therefore, for negative binomial models, K was allowed to fluctuate with the model that had the lowest AICc score to examine when the population estimate levels off, as well as changes in p, and goodness of fit.

### Univariate Relationships and Habitat Associations

Present within the system are species of at least nine different genera of Elmidae. Most of these species are likely more riverine, however, our interest was in spring-adapted species. To examine community structure of elmid beetles and determine spring-adapted species present, lures were set in a longitudinal fashion from the origin of the spring to parts of the spring-run more influenced by external temperature and mixing of the spring water with organic material. A total of 17 other lures were set out during the course of the study (March, April, and November) downstream of the spring origin within the spring-run, which we termed the transition zone. The transition zone has subtle changes in water chemistry compared to the spring origin which may restrict the distribution of spring-adapted or associated organisms. Lures were set in the same fashion as the abundance model lures and left in place for the same amount of time. When lures were collected, elmid adults and larvae were counted and returned to the site of capture.

To examine differences between the two elmid communities present at the spring origin and the transition zone, and to determine which elmids appear to be spring adapted, indicator species analysis [35] was conducted using the statistical package “labdsv” in R [36]. Count data from the abundance model was averaged (n = 32) from the four events and compared to the transition zone lure averages (n = 17). Other univariate relationships with water chemistry and substrate were explored using Pearson correlation analysis. Variables that were correlated with count data at ± 0.50 were examined using linear regression models. This was done by averaging the count data from all events and testing against the data collected during the mapping event (i.e. specific conductance, temperature, etc). To test for correlations with the substrate, the designation of substrate types was expanded to run from 0 to15 using a modified Wentworth scale.

## Results

### N-mixture Abundance Model

The final data set consisted of 32 sites, 28 spring sites and four seep sites. Some seep sites were disregarded due to the low flow and the collection method used. Specifically, most seep sites were not conducive to the lure method and alternate spring sites were used instead. Final sites used in the abundance models are presented in Table 1. A total of 3,122 adult *H. cf. glabra* were counted from lures during the four events. The highest capture rate occurred in November (n = 1,078), while the lowest capture rate (n = 602) was in August. Finegan Springs consistently had higher capture rates than at Dolan Springs.

**Table 1.** Randomly selected spring sites in Val Verde County used to populate *Heterelmis c.f. glabra* abundance models.

| Springs | Site | Type | Flow | Substrate | Temperature | Conductivity | DO | pH | TDS |
| --- | --- | --- | --- | --- | --- | --- | --- | --- | --- |
| BH | 8 | group of springs | 2 | 3 | 22.11 | 495 | 7.63 | 6.82 | 0.3175 |
| BH | 9 | group of springs | 2 | 3 | 22.12 | 490 | 7.97 | 7.06 | 0.3139 |
| BH | 1 | orifice | 3 | 4 | 22.13 | 495 | 7.72 | 6.91 | 0.3169 |
| BH | 2 | orifice | 4 | 4 | 21.98 | 495 | 7.36 | 7.60 | 0.3168 |
| BH | 5 | orifice | 2 | 3 | 22.03 | 537 | 7.60 | 7.13 | 0.3435 |
| FS | 84.0 | orifice | 2 | 4 | 22.34 | 498 | 8.14 | 7.14 | 0.3188 |
| FS | 94.7 | Seep | 1 | 2 | 21.81 | 498 | 8.68 | 7.37 | 0.3194 |
| FS | 105.0 | orifice | 2 | 3 | 22.45 | 497 | 8.09 | 7.34 | 0.3180 |
| FS | 110.0 | orifice | 2 | 3 | 22.43 | 489 | 8.24 | 7.22 | 0.3134 |
| FS | 151.5 | orifice | 3 | 3 | 22.48 | 497 | 8.13 | 7.16 | 0.3183 |
| FS | 175.7 | orifice | 4 | 2 | 22.49 | 498 | 8.12 | 7.20 | 0.3185 |
| FS | 184.3 | orifice | 4 | 4 | 22.48 | 498 | 7.88 | 7.16 | 0.3185 |
| FS | 191.0 | orifice | 3 | 6 | 22.48 | 497 | 8.03 | 7.19 | 0.3178 |
| FS | 212.0 | Seep | 1 | 6 | NA | NA | NA | NA | NA |
| FS | 280.5 | group of springs | 3 | 2 | 22.50 | 498 | 8.04 | 7.19 | 0.3185 |
| FS | 285.0 | group of springs | 3 | 6 | 22.50 | 497 | 7.96 | 7.20 | 0.3186 |
| FS | 310.4 | group of springs | 3 | 4 | 22.49 | 498 | 7.67 | 7.28 | 0.3186 |
| FS | 319.0 | orifice | 3 | 6 | 22.49 | 497 | 8.10 | 7.20 | 0.3179 |
| FS | 291.5 | orifice | 5 | 6 | 22.49 | 498 | 8.13 | 7.22 | 0.3188 |
| DC | 43.0 | group of springs | 3 | 4 | 22.29 | 477 | 7.27 | 7.09 | 0.3053 |
| DC | 120.0 | Seep | 1 | 6 | NA | NA | NA | NA | NA |
| DC | 135.3 | orifice | 3 | 1 | 21.86 | 485 | 6.75 | 7.15 | 0.3191 |
| DC | 252.7 | orifice | 4 | 3 | 22.56 | 507 | 8.08 | 7.05 | 0.3322 |
| DC | 261.5 | group of springs | 2 | 3 | 22.49 | 497 | 6.73 | 7.24 | 0.3301 |
| DC | 264.5 | orifice | 3 | 3 | 22.42 | 485 | 7.53 | 7.16 | 0.3107 |
| DC | 267.5 | Seep | 1 | 1 | NA | NA | NA | NA | NA |
| DC | 278.6 | orifice | 3 | 3 | 22.55 | 486 | 7.77 | 7.14 | 0.3114 |
| DC | 279.7 | orifice | 3 | 3 | 22.48 | 494 | 7.55 | 7.21 | 0.3250 |
| DC | 641.0 | orifice | 3 | 3 | 22.54 | 479 | 8.16 | 7.27 | 0.3069 |
| DC | 644.4 | orifice | 3 | 3 | 22.53 | 479 | 8.05 | 7.13 | 0.3071 |
| DC | 650.5 | group of springs | 2 | 6 | NA | NA | NA | NA | NA |
| DC | 658.2 | group of springs | 3 | 6 | 21.70 | 488 | 8.12 | 7.45 | 0.3169 |

BH = Blue Hole; FS = Finegan Springs; DC = Dolan Creek; NA = too shallow to sample.

The top ranking abundance models for both the Poisson and NB models contained the additive effects of rain, site and flow within the detection parameter (global models). The global negative binomial model had the lowest AIC_c_ value with an overall AIC_c_ weight of 0.84 compared to the other 12 models. There was a change in AICc of 3 compared to the following model and a separation of AIC_c_ of 424 to the closest Poisson model (Table 2). All NB models had lower AIC_c_ scores than the Poisson. The global Poisson model scored 143 AIC_c_ points lower than the null negative binomial model. Site and flow had positive relationships within the models with rain having a negative effect on the detection of riffle beetles.

**Table 2.**
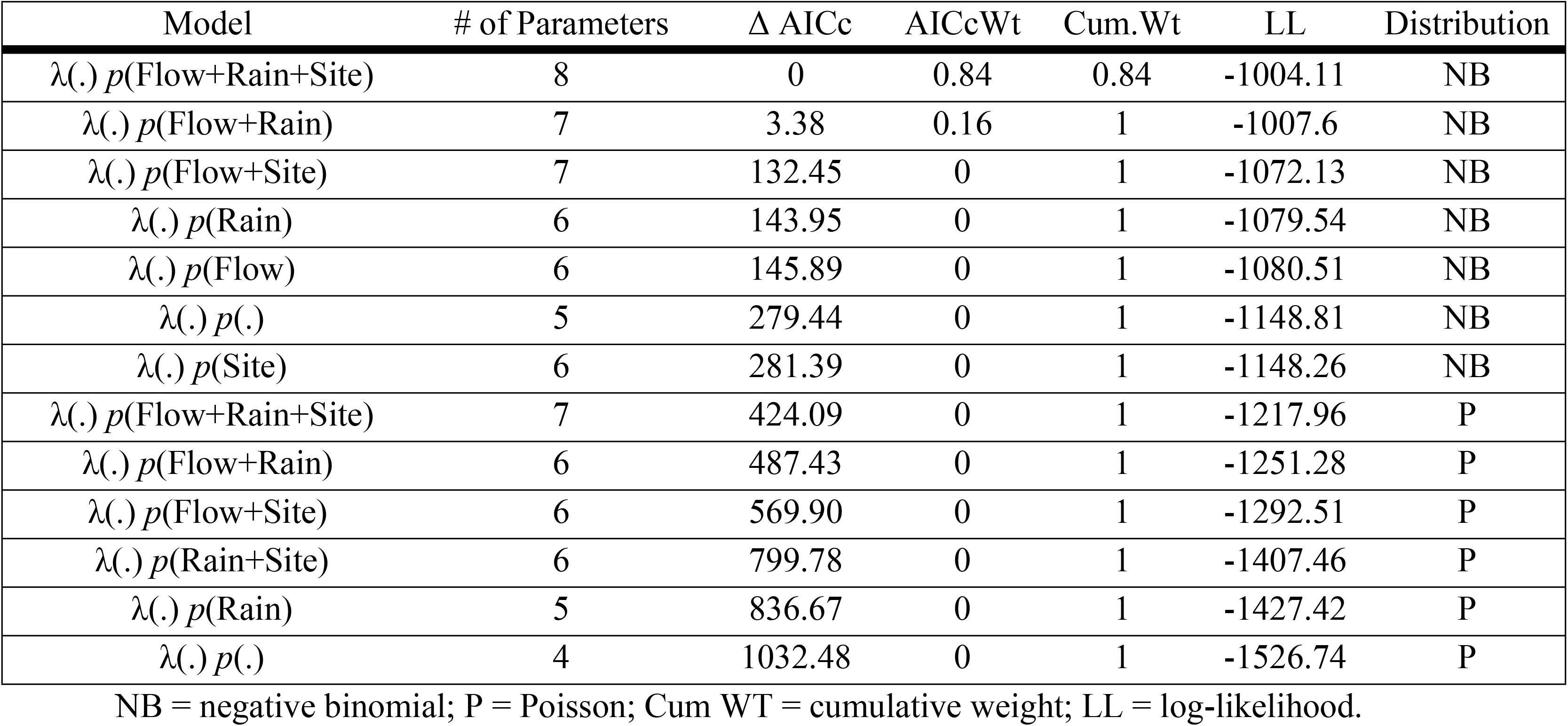
Results from negative binomial (NB) and Poisson (P) open season N-mixture abundance models for *Heterelmis c.f. glabra*.

| Model | # of Parameters | $\Delta$ AICc | AICcWt | Cum. Wt | LL | Distribution |
| --- | --- | --- | --- | --- | --- | --- |
| $\lambda(.) p(\text{Flow}+\text{Rain}+\text{Site})$ | 8 | 0 | 0.84 | 0.84 | -1004.11 | NB |
| $\lambda(.) p(\text{Flow}+\text{Rain})$ | 7 | 3.38 | 0.16 | 1 | -1007.6 | NB |
| $\lambda(.) p(\text{Flow}+\text{Site})$ | 7 | 132.45 | 0 | 1 | -1072.13 | NB |
| $\lambda(.) p(\text{Rain})$ | 6 | 143.95 | 0 | 1 | -1079.54 | NB |
| $\lambda(.) p(\text{Flow})$ | 6 | 145.89 | 0 | 1 | -1080.51 | NB |
| $\lambda(.) p(.)$ | 5 | 279.44 | 0 | 1 | -1148.81 | NB |
| $\lambda(.) p(\text{Site})$ | 6 | 281.39 | 0 | 1 | -1148.26 | NB |
| $\lambda(.) p(\text{Flow}+\text{Rain}+\text{Site})$ | 7 | 424.09 | 0 | 1 | -1217.96 | P |
| $\lambda(.) p(\text{Flow}+\text{Rain})$ | 6 | 487.43 | 0 | 1 | -1251.28 | P |
| $\lambda(.) p(\text{Flow}+\text{Site})$ | 6 | 569.90 | 0 | 1 | -1292.51 | P |
| $\lambda(.) p(\text{Rain}+\text{Site})$ | 6 | 799.78 | 0 | 1 | -1407.46 | P |
| $\lambda(.) p(\text{Rain})$ | 5 | 836.67 | 0 | 1 | -1427.42 | P |
| $\lambda(.) p(.)$ | 4 | 1032.48 | 0 | 1 | -1526.74 | P |

NB = negative binomial; P = Poisson; Cum WT = cumulative weight; LL = log-likelihood.

Model selection was based upon the AICc score initially. Subsequently, the preliminary selection of the global NB at a K of 400 was modeled with a goodness of fit and evaluated at different levels of K. The levels of K ranged from the default value of 157 to the selected value of 600 (Fig 3). The abundance parameter estimates of K at 500 and 600 became inseparable. However, the probabilities of detection decreased as K level increased. Goodness of fit tests at 500 K showed the mathematical fit of the model (p = 0.08) with a c-hat (ĉ) of 1.68 (Table 3). Goodness of fit tests for the 600 K model showed a lack of fit (p = 0.01) and had a higher ĉ (1.91) than the 500 K model. Therefore, the global NB model at a K of 500 was selected as the appropriate model for this data set.

**Fig 3.**
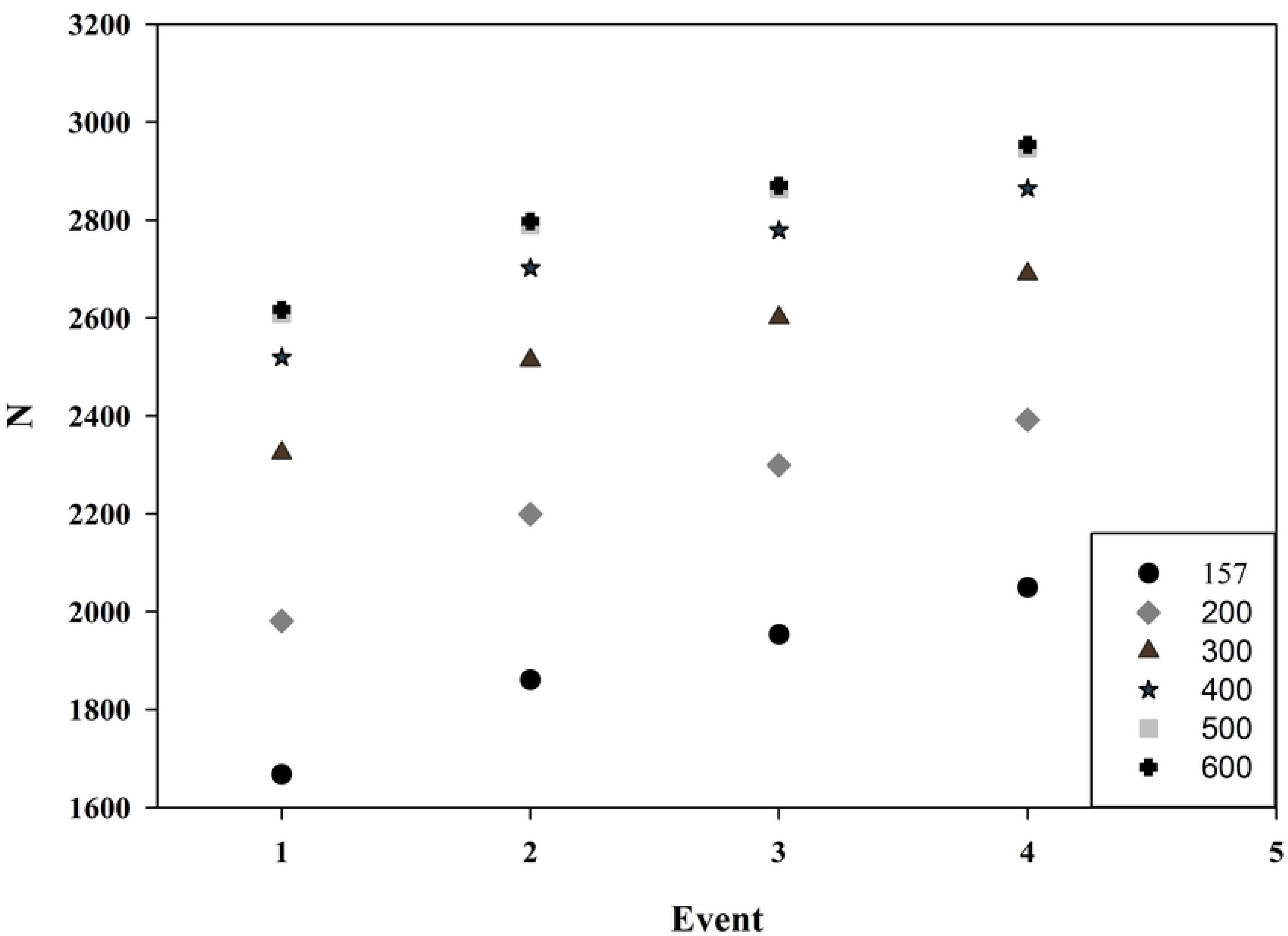
Results from the selected N-mixture model. Negative binomial mixture of global models showing range of site level estimates with varying values of K.

**Table 3.** Selected global abundance model and the relationship with K.

| Model | K | AICc | GOF | $\hat{c}$ |
| --- | --- | --- | --- | --- |
| $\lambda(.) p(\text{Flow}+\text{Rain}+\text{Site})$ | 600 | 2030.13 | 0.01 | 1.91 |
| $\lambda(.) p(\text{Flow}+\text{Rain}+\text{Site})$ | 500 | 2030.14 | 0.08 | 1.68 |
| $\lambda(.) p(\text{Flow}+\text{Rain}+\text{Site})$ | 400 | 2030.48 | 0.03 | 1.89 |
| $\lambda(.) p(\text{Flow}+\text{Rain}+\text{Site})$ | 300 | 2032.62 | NA | NA |
| $\lambda(.) p(\text{Flow}+\text{Rain}+\text{Site})$ | 200 | 2040.48 | NA | NA |
| $\lambda(.) p(\text{Flow}+\text{Rain}+\text{Site})$ | 157 | 2052.59 | 0.02* | 2.1* |

K = values for the upper bound of integration.
\*Asterisks indicate warnings associated with the analysis

The abundance parameter estimates taken from the global NB model at 500 K for combined spring sites sampled increased over the year from 2,609 (+690/-874) to 2,946 (+451/-479). Overall probabilities of detection in the NB 500 model ranged from 0.07 to 0.80 on average (Table 4). Site 291 at Finegan Springs had the highest probability of detection for the data set (0.856). Site 120 at Dolan Springs had the lowest probability of detection (0.045). The results suggest that as the flow increases the probability of detecting a beetle increases. The probability of detection is the lowest for the events (April and August) with rainfall during the deployment.

**Table 4.** Probabilities of detection by site along with averages calculated from the selected negative binomial global model.

| Spring | Site | March | May | August | November | Average |
| --- | --- | --- | --- | --- | --- | --- |
| Blue | 8 | 0.241 | 0.134 | 0.134 | 0.241 | 0.187 |
| Blue | 9 | 0.241 | 0.134 | 0.134 | 0.241 | 0.187 |
| Blue | 1 | 0.457 | 0.291 | 0.291 | 0.457 | 0.374 |
| Blue | 2 | 0.691 | 0.522 | 0.522 | 0.691 | 0.607 |
| Blue | 5 | 0.241 | 0.134 | 0.134 | 0.241 | 0.187 |
| Finegan | 84.0 | 0.241 | 0.134 | 0.134 | 0.241 | 0.187 |
| Finegan | 94.7 | 0.107 | 0.055 | 0.055 | 0.107 | 0.081 |
| Finegan | 105.0 | 0.241 | 0.134 | 0.134 | 0.241 | 0.187 |
| Finegan | 110.0 | 0.241 | 0.134 | 0.134 | 0.241 | 0.187 |
| Finegan | 151.5 | 0.457 | 0.291 | 0.291 | 0.457 | 0.374 |
| Finegan | 175.7 | 0.691 | 0.522 | 0.522 | 0.691 | 0.607 |
| Finegan | 184.3 | 0.691 | 0.522 | 0.522 | 0.691 | 0.607 |
| Finegan | 191.0 | 0.457 | 0.291 | 0.291 | 0.457 | 0.374 |
| Finegan | 212.0 | 0.107 | 0.055 | 0.055 | 0.107 | 0.081 |
| Finegan | 280.5 | 0.457 | 0.291 | 0.291 | 0.457 | 0.374 |
| Finegan | 285.0 | 0.457 | 0.291 | 0.291 | 0.457 | 0.374 |
| Finegan | 310.4 | 0.457 | 0.291 | 0.291 | 0.457 | 0.374 |
| Finegan | 319 | 0.457 | 0.291 | 0.291 | 0.457 | 0.374 |
| Finegan | 291.5 | 0.856 | 0.744 | 0.744 | 0.856 | 0.800 |
| Dolan | 43.0 | 0.403 | 0.248 | 0.248 | 0.403 | 0.326 |
| Dolan | 120.0 | 0.087 | 0.045 | 0.045 | 0.087 | 0.066 |
| Dolan | 135.3 | 0.403 | 0.248 | 0.248 | 0.403 | 0.326 |
| Dolan | 252.7 | 0.643 | 0.467 | 0.467 | 0.643 | 0.555 |
| Dolan | 261.5 | 0.203 | 0.110 | 0.110 | 0.203 | 0.156 |
| Dolan | 264.5 | 0.403 | 0.248 | 0.248 | 0.403 | 0.326 |
| Dolan | 267.5 | 0.087 | 0.045 | 0.045 | 0.087 | 0.066 |
| Dolan | 278.6 | 0.403 | 0.248 | 0.248 | 0.403 | 0.326 |
| Dolan | 279.7 | 0.403 | 0.248 | 0.248 | 0.403 | 0.326 |
| Dolan | 641.0 | 0.403 | 0.248 | 0.248 | 0.403 | 0.326 |
| Dolan | 644.4 | 0.403 | 0.248 | 0.248 | 0.403 | 0.326 |
| Dolan | 650.5 | 0.203 | 0.110 | 0.110 | 0.203 | 0.156 |
| Dolan | 658.2 | 0.403 | 0.248 | 0.248 | 0.403 | 0.326 |

### Univariate Relationships and Habitat Associations

Indicator analysis on the Elmidae communities collected on transition lures and spring origin lures showed that *H. cf. glabra* adults and larvae are associated with spring sources (p = 0.001; Table 5). The associations of *Microcylloepus* sp. adults and larvae with spring sites were not significant although were shown to associate with the transition zone more than the spring origin. *Phanocerus* larvae were significantly associated with the spring origin sites (p = 0.001). Other genera were not detected within the transition zone or the spring area.

**Table 5.**
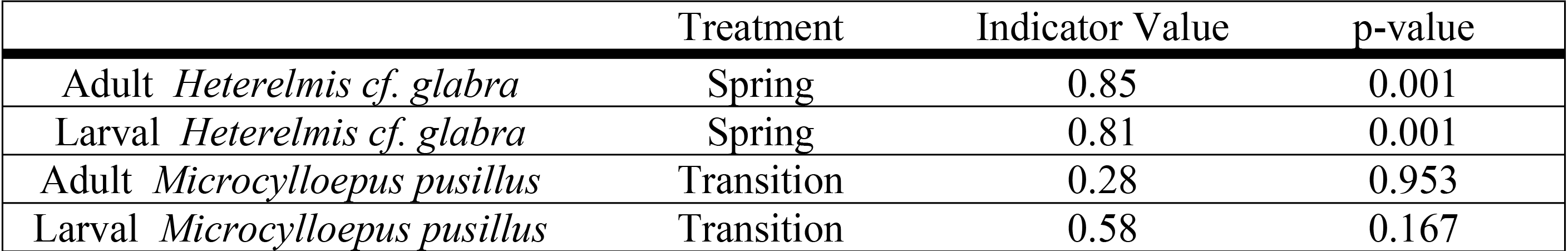
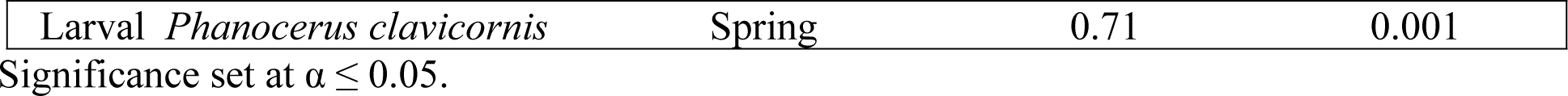
Indicator species analysis results showing treatment, relevant indicator value, and significance.

|  | Treatment | Indicator Value | p-value |
| --- | --- | --- | --- |
| Adult *Heterelmis cf. glabra* | Spring | 0.85 | 0.001 |
| Larval *Heterelmis cf. glabra* | Spring | 0.81 | 0.001 |
| Adult *Microcylloepus pusillus* | Transition | 0.28 | 0.953 |
| Larval *Microcylloepus pusillus* | Transition | 0.58 | 0.167 |
| Larval *Phanocerus clavicornis* | Spring | 0.71 | 0.001 |

Significance set at $\alpha \leq 0.05$ .

Only three abiotic variables (temperature, flow, and substrate) had significant correlations with either adult or larval *H. cf. glabra* average count data. The first relationship was between the measured temperature at the time of the mapping and the average count data of adult *H. cf. glabra* (Fig 4). As the temperature increases there is a significant increase in the presence of adult *H. cf. glabra* (F_1,26_ = 10.14, r^2^ = 0.28, p = 0.003). *Heterelmis cf. glabra* also exhibited a significant relationship with flow (Fig 4). As flow increased the average number of adult *H. cf. glabra* collected increased (F_1,30_ = 16.64, r^2^ = 0.35, p = 0.003). A negative relationship was observed with *H. cf. glabra* larvae and substrate (Fig 4). As substrate size increased the average count of *H. cf. glabra* larvae decreased (F_1,30_ = 9.39, r^2^ = 0.23, p = 0.004), suggesting potential habitat partitioning between adult and larval *H. cf. glabra* as the adult correlation was positive although not significant.

**Figure 4.**
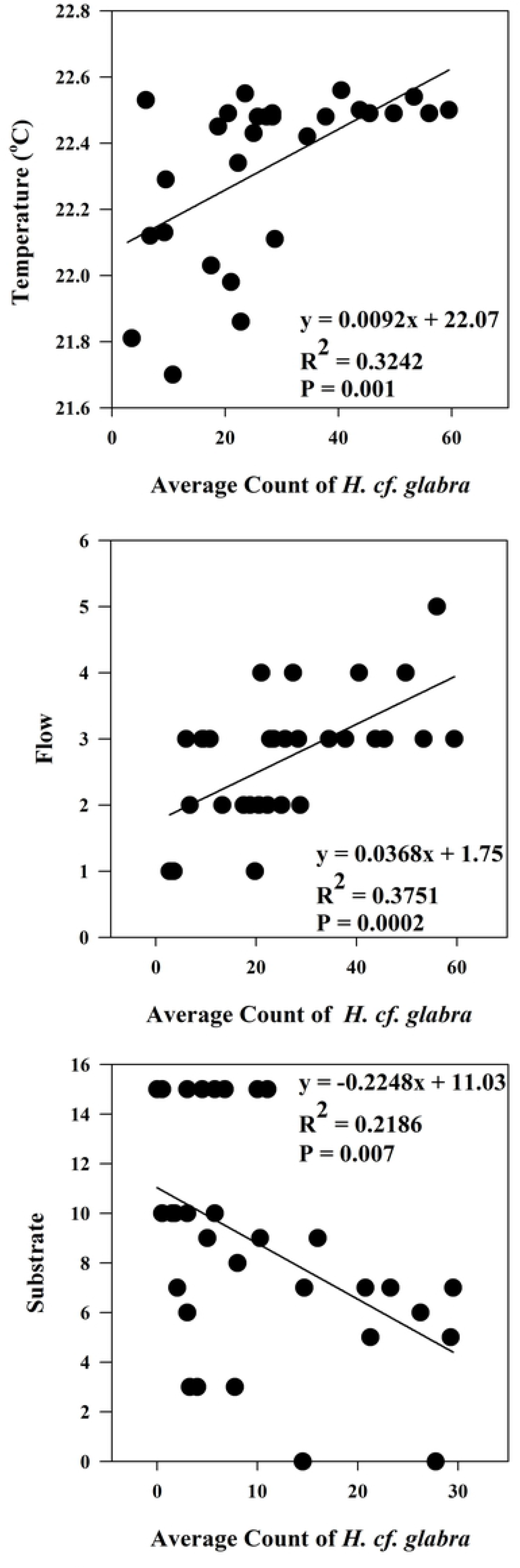
Univariate relationships with measured abiotic parameters of adult and larval *Heterelmis cf. glabra* collected in Val Verde, TX. Presented are relationships for temperature, flow and substrate. Significance set at α ≤ 0.05

## Discussion

The results from the models suggest that rainfall, flow and the site were important to detections of *H. cf. glabra* within this area of its distribution. Individually, the covariates had different levels of fit to the models. Flow and site both had significant positive relationships highlighting differences between the site level estimates of abundance at different flows and between the sites. Rainfall did have a negative relationship with the probability of detection, which was hypothesized to have such an effect due to the sedimentation issues and local changes in water chemistry during and post rainfall.

The calculated estimates from the NB global model of site level abundance and total abundance seem realistic and ecologically plausible. Although the area from which the lure is sampling the beetles is not known, the sampling consistency at all sites provides reliable comparisons between sites. As mentioned previously, flow displayed a strong positive relationship to the probability of detecting beetles. The flow at each site may in part determine the area from which these beetles were drawn. Therefore, the greater the flow the more potential microhabitat from which to draw beetles to the lure.

When abundance parameter estimates from model runs are compared to raw data, the sites with lower raw counts seem to have higher predicted abundance than in the calculated data for sites with higher raw counts. For example, seep sites, ranked with a flow of ‘one’, have the lowest probabilities of detection among all of the flow categories and higher estimates of abundance than the count data for these sites. The model appears to be accounting for riffle beetles potentially not present at the site due to the low probability of detection at these lower flowing sites, suggesting a sampling issue with the seep sites. Therefore, one scenario would be where the beetles are not being detected, although present, thereby inflating the abundance score associated with these types of sites. Another possibility is that at these lower flowing spring sites, the zeros in count data could be true zeros not modeled within the predicted data set. Either scenario discussed above, is highlighting the need for a better estimate of flow than the categorical type that was used for these models, or disregarding seep sites for these types of models. Adding flow as a continuous variable, collected at deployment and pick up, may provide more reliable site level abundances than using the ordinal flow designation. Monitoring the flow at each site with a weir or other technique at the beginning and end of each event may provide the subtle changes in flow data that could be used as a sampling covariate to explain fluctuations in abundance over time.

The negative binomial models were ranked lower by AICc scores than any of the Poisson models. This seems to be the trend for the NB models showing higher site level abundance when compared to the corresponding Poisson models. In some cases, NB models in other studies have produced approximately double the abundance of the normal territory mapping method [24; Add]. Goodness of fit test on selected NB model results I this study showed acceptable fit as the Poisson models were not in congruence [24]. Subsequently, a recent study [37] suggested the estimates from K’ery et al. (2005; [24]) may be more reliable had the Poisson distribution been selected and not the NB model. For this study, the Poisson models (global and null) both had very high values (+ 15) of ĉ, indicating lack of fit within the model. In this case, based upon the fitted levels of K, goodness of fit tests, and the AICc score the NB model at 500 K was selected.

There are a number of differences between our study organism and the organisms studied in the available literature where n-mixture models were employed. For example, our site level population estimates are consistently larger than many other reported estimates of lambda [38, 39]. Due to the small size of the beetle there is the potential to have large numbers of individuals detected within or on the lure. Many studies have much shorter sampling periods (e.g., five to ten minutes for bird surveys; [13, 14], however, with this study the lures were deployed for weeks as it takes time for biofilms to grow on the lures attracting the beetles. Although N-mixture models have not been used for many studies involving invertebrates, this approach is been useful with this particular species.

Flow and temperature were both significant abiotic variables when compared to the average adult *H. cf. glabra* count data. To determine if flow is the actual mechanism or if size of the spring-run is creating larger abundances, size of the run should be considered a site covariate. Greater spring flows presumably sustain more suitable habitat from which to draw beetles to the lure. Temperature values were collected during the mapping event in January of 2016, therefore, lower temperatures may show spring sites with more exposure to the environment or shallow laminar flow which is more susceptible to temperature fluctuations at the surface or farther away from the origin of the spring. Future efforts may consider incorporating temperature as a covariate measured at the beginning and end of each sampling event.

Overall, N-mixture models have great potential as a monitoring tool for rare, small, and difficult to collect interstitial species, such as riffle beetles. In order to determine trends within the population, regular monitoring of the beetles should be done with set monitoring locations at least three times a year for a number of years. In addition, sites should be added if possible to increase the sample size in order to conduct hypothesis tests on covariates that influence abundance, not only the probability of detection. After these baseline surveys are conducted, future surveys could be compared and changes in surface populations and available habitat of spring-adapted riffle beetles could be elucidated.

The Devils River has long been recognized as a least disturbed stream and has many unique species associated with its watershed. While anthropogenic stressors have been lacking in the area historically, advances in nontraditional oil and gas activity has created opportunity for industrial expansion in this region. Over 47,000 oil and gas wells were permitted in the Permian Basin region between 2011 and 2016, with water usage per well increasing 770% (to up to 42,500 m^3^ per well) during that same period [40]. Current commercial estimates predict continued growth of production within the region over the next few years, suggesting the demand for water in the region will increase. Groundwater dependent rivers and streams, such as the Devils River, may experience decreases in springflow and thus overall streamflow due to pumping activity. Modeling demonstrated that “production of groundwater in the [Devils River] basin will result in a proportional reduction in the flow of the Devils River” with the impact “most pronounced during low flow conditions” potentially impacting the ecology of the system as spring discharge points in the river are extinguished [1]. While H*. cf glabra* currently has no protected status, the need for such protections would grow if known populations are negatively impacted by water development. Incorporating springflow metrics protective of spring habitats into groundwater management is needed to reduce potential impacts to rare spring dependent species, such as *H. cf glabra* while ensure sustainable water supplies.

## Acknowledgments

We would like to thank The Nature Conservancy (namely Deirdre Hisler, John Karges, and Ryan Smith) for access to the habitat for this project. We also thank Texas Parks and Wildlife Department (namely Joe Joplin and David Riskind) for access to the habitat. Additionally, we thank anonymous reviewers, the staff of U. S. Fish and Wildlife Service (Texas Fish and Wildlife Conservation Office, San Marcos Aquatic Resources Center, and Ecological Services), River Studies (Texas Parks and Wildlife Department), and Parvathi Nair (Texas State University) for help in the field. The views presented herein are those of the authors and do not necessarily represent those of the U.S. Fish and Wildlife Service or Texas Parks and Wildlife Department.

